# Plausible Mechanistic Insights in Biofilm Eradication Potential of against *Candida* spp. using In Situ Synthesized Tyrosol Functionalized Chitosan Gold Nanoparticles as a versatile Antifouling Coating on Implant Surfaces

**DOI:** 10.1101/2021.09.30.462644

**Authors:** Tara Chand Yadav, Payal Gupta, Saakshi Saini, Vikas Pruthi, Ramasare Prasad

**Affiliations:** Department of Biosciences and Bioengineering, Indian Institute of Technology Roorkee, Roorkee – 247667, Uttarakhand, India

**Keywords:** Antibiofilm, *Candida*, chitosan, gold nanoparticles, tyrosol, transcriptional analysis

## Abstract

In the present study, tyrosol functionalized chitosan gold nanoparticles (Chi-TY-AuNP’s) were prepared as an alternative treatment strategy to combat fungal infections. Various biophysical techniques were used to characterize the synthesized Chi-TY-AuNP’s. The antifungal and antibiofilm activities of Chi-TY-AuNP’s were evaluated against *C. albicans* and *C. glabrata* and efforts have been made to elucidate the possible mechanism of action. Chi-TY-AuNP’s showed a high fungicidal effect against both sessile and planktonic cells of *Candida* spp. Additionally, Chi-TY-AuNP’s completely eradicated (100%) the mature biofilms of both the *Candida* spp. FESEM analysis highlighted the morphological alterations in Chi-TY-AuNP’s treated *Candida* biofilm cells. Effect of Chi-TY-AuNP’s on the ECM components showed significant reduction in protein content in *C. glabrata* biofilm and substantial decrease in extracellular DNA (eDNA) content of both the *Candida* spp. ROS generation analysis using DCFDA-PI staining showed high ROS levels in both the *Candida* spp., whereas pronounced ROS production was observed in Chi-TY-AuNP’s treated *C. glabrata* biofilm. Biochemical analysis revealed decreased ergosterol content in Chi-TY-AuNP’s treated *C. glabrata* cells, while inconsequential changes were observed in *C. albicans*. Furthermore, the transcriptional expression of selected genes (ergosterol biosynthesis, efflux, sterol importer, and glucan biogenesis) was reduced in *C. glabrata* in response to Chi-TY-AuNP’s except *ERG11* and *CDR1*. Conclusively the result showed the biofilm inhibition and biofilm eradication efficacy of Chi-TY-AuNP’s in both the *Candida* spp. Findings of the present study manifest Chi-TY-AuNP’s as a potential therapeutic solution to *Candida* biofilm-related chronic infections and overcome biofilm antifungal resistance.

## INTRODUCTION

Candidiasis is amongst the utmost prevalent nosocomial pathogenic infections triggered by *Candida*, primarily in immune-compromised patients with an approximately 40% mortality rate.^1^ *Candida* is a commensal polymorphic yeast responsible for superficial skin infections to deep tissue invasions. *Candida albicans* is the most ubiquitous and menacing pathogen, followed by non-albicans *Candida* spp., where *C. glabrata* and *C. tropicalis* stands out as the most prevalent spp.^2^ Epidemiological reports from North America suggest that there are frequent incidences of candidiasis due to *C. glabrata*, also prevalent in the Asia-pacific region.^3^ The virulence properties of *Candia* spp. are attributed to its capability to form biofilm-a sessile, multicellular community where microbes encapsulate themselves in a self-secreted extracellular matrix (ECM).^4^ This ECM is a conglomeration of biomolecules and hydrolytic enzymes, specifically lipase, proteinase and phospholipase. Besides biofilm, adhesin proteins and persister cells further boost their resistant nature and contribute to recurrent candidiasis.^5^ Persister cells belong to the biofilm community exhibiting lower growth rates, active drug efflux, high resistance to antimicrobial treatment and entirely different from free-floating planktonic cells.^6^ These biochemical events are attributed to biofilm growth and act cohesively to combat the antifungals therapeutic action.

Inside biofilm, cells communicate with each other harmonically, and this cellular signaling is known as quorum sensing (QS), medicated by quorum sensing molecules (QSMs). Tyrosol (2-(4-hydroxyphenyl)ethanol) is a QSMs molecule produced by *C. albicans* that stimulates the formation of germ tubes and initiates the development of hyphae to facilitate biofilm formation, whereas its exogenous administration acts antagonistically.^7, 8^ The antibacterial and antifungal properties of tyrosol (TY) have been substantially explored in recent years; however, these investigations unravel insufficient mechanistic insight into tyrosol’s mode of action in *Candida*.^9, 10^ Increasing incidences of *Candida-*related nosocomial infections and emerging multidrug resistance (MDR) *Candida* spp. are a global concern due to the high mortality rate. Rapidly increasing resistance to major antifungals facilitated through multiple mechanisms limited the treatment options for *Candida*-related infections.^11^ Therefore, the development of novel antifungals, combinatorial therapies and the quest for alternative therapeutic strategies are crucial to mitigate the recalcitrant biofilms and repercussions of rapidly growing antimicrobial resistance.

In recent years, nanoparticles emerged as an invincible tool in intracellular delivery of antimicrobials due to their minimal size, high surface to volume ratio, enhanced cellular uptake and sub-cellular drug retention properties.^12^ Among various metallic nanoparticles, gold in nanoparticulated form has shown immense potential in antimicrobial drug delivery owing to their high therapeutic efficacy against bacteria and fungi followed by low cytotoxicity and high biocompatibility against mammalian cells.^13^ Chitosan is a natural cationic polysaccharide comprising of glucosamine and N-acetyl glucosamine units. The presence of different functional groups such as hydroxyl (–OH), amine (–NH_2_) and carbonyl (> C=O) renders chitosan an extensively used biopolymer for the synthesis of colloidal nanoparticles acting as a reducing and stabilizing agent.^14^ The broad-spectrum antimicrobial action of chitosan against a wide variety of microbes is attributed to its polycationic nature contributed by NH_3_^+^ groups of glucosamine units. The cationic tail of chitosan interacts with the negatively charged cytoplasmic membrane of microbes via electrostatic interactions leading to extensive cell surface modifications. Subsequently, this results in cellular internalization of nanoparticles, leakage of intracellular contents, ultimately resulting in inhibition of DNA transcription as well as RNA and protein synthesis.^15^ Besides, chitosan’s toxicity towards bacterial cells and mammalian cells safety has ruled out its biocompatibility issues.^16^

A combinatorial drug delivery system that enables easy transport of drugs to the target site is a potential strategy in treating *Candida* infections.^17^ Commonly used antifungals such as azoles and polyenes etc., can be metamorphosed into nanoparticles to improve antifungal agents’ efficacy compared to the conventional therapeutic regimen.^12^ In view of the exceptional physicochemical characteristics of chitosan and gold renders them as excellent biomaterials in the fabrication of effective nanoformulation. Besides, chitosan’s biocompatibility and gold being inert along with antimicrobial efficacy offer a promising horizon in the development of noncytotoxic chitosan-gold nanoparticles.^18^ In addition, chitosan-based nanoformulations possess a high positive surface charge, which helps in the transportation of molecules/ drugs across the cell membrane, high drug payload and enhanced cellular uptake. Such carrier systems would enhance the cellular internalization of active molecule/ drug due to the electrostatic interaction between nanoparticles and cell membrane leading to enhanced penetration in *Candida* biofilm and its membrane disruption, thus providing a smart strategy to combat the menacing effect of biofilm. The all-pervading chitosan-gold nanoparticle’s ability inside the biofilm niche is very high; therefore, they possess high antimicrobial activity, which is crucial for combating *Candida* spp. mediated superficial or systemic infections.

In the present study, TY functionalized chitosan gold nanoparticles (Chi-TY-AuNP’s) were synthesized by *in situ* facile method to harness synergistic effect via targeting both the fungal cells and biofilm matrix. The antifungal and antibiofilm potency of synthesized Chi-TY-AuNP’s was investigated against *C. albicans* and *C. glabrata*. Further, efforts have been made to elucidate the mode of action of Chi-TY-AuNP’s by examining their impact on ROS generation, cell surface hydrophobicity, ECM composition and membrane ergosterol content in biofilms of both the *Candida* spp., besides, transcriptional expression of selected *C. glabrata* genes were also evaluated.

## MATERIALS AND METHODS

### Chemicals and Reagents

Chitosan (low molecular weight), glacial acetic acid, and gold (III) chloride trihydrate (HAuCl_4_•3H_2_O) (≥ 99.9%) were procured from Sigma-Aldrich, St. Louis, USA. Tyrosol (purity > 98.0%) was purchased from TCI Chemicals (India), Pvt. Ltd. All media components were procured from Himedia, India, BCA protein assay kit (Sigma-Aldrich, USA), RNeasy kit (Qiagen, Germany), Verso cDNA synthesis kit (Thermo Fisher Scientific, USA) and other chemicals were obtained from Sigma-Aldrich, USA. All glassware was treated with Aqua regia and rinsed with Milli-Q water, before proceeding with the experiments the glassware was dried in the hot-air oven for 5 h.

### Synthesis of Chi-TY-AuNP’s

Chitosan flakes were dissolved in 100 mL of Milli-Q water in 1% CH_3_COOH to prepare 0.2% (w/v) chitosan solution, followed by the addition of 91.9 μ (136 mM) of gold (III) chloride trihydrate (HAuCl_4_•3H_2_O). Subsequently, it was heated for 15 min at 90°C in a heating mantle, accompanied by continuous stirring. The change in color of the mixture from colorless to ruby-red indicated the formation of Chi-AuNP’s. For drug loading, Chi-AuNP’s were stirred magnetically with 1 mg/mL of TY for 24 h at 25°C and further incubated at 4°C for 48 h. The mixture was then subjected to ultracentrifugation for 30 min at 30,000 rpm, and Milli-Q water was used for pellet redispersion.^19, 20^

### Characterization of Synthesized Chi-TY-AuNP’s

Shimadzu-1700 UV–visible spectrophotometer (resolution 1 nm; scanning λ = 400-800 nm) was used to determine the surface plasmon resonance (SPR) of Chi-TY-AuNP’s. ’Image J 1.49’ software (National Institute Health, USA) was used to calculate the distribution of Chi-TY-AuNP’s diameter. The polydispersity index (PDI), dynamic light scattering (DLS) (hydrodynamic size), and surface charge of Chi-TY-AuNP’s were estimated with Zetasizer (Malvern Zetasizer Nano ZS90, UK) at 25°C. High-resolution transmission electron microscopy (HRTEM, FEI Tecnai G2, USA), operating at the voltage of 200 kV, was utilized to determine the morphology of Chi-TY-AuNP’s. For atomic force microscopy (AFM, AFM-STM, Ntegra T-150, Ireland) analysis, Chi-TY-AuNP’s were diluted (10 times) with Milli-Q water and dried out onto a clean glass slide under vacuum at 25°C for 24 h. Further, dried Chi-TY-AuNP’s were examined for morphological analysis.

### Fourier-Transform Infrared (FTIR) Analysis

The functional group characterization of Chi-AuNP’s and Chi-TY-AuNP’s, along with TY and chitosan, were performed by FTIR spectroscopy. Vibrational frequencies in the infra-red (IR) region (4000-400 cm^−1^) were analyzed by the KBr pellet method using a Thermo Nicolet spectrometer, determining the formation and chemical modifications of Chi-TY-AuNP’s.

### Tyrosol Loading Efficiency of Chi-TY-AuNP’s

After the synthesis of Chi-TY-AuNP’s, the unreacted drug and other substrates were removed by continuous dialysis in a 14 kD cut-off membrane for 72 h. A known volume of aqueous dispersed Chi-TY-AuNP’s was subjected to probe sonication for 15 min, provided with 2 s “On” and 3 s “Off” pulse duration. Using UV-visible spectrophotometer, the amount of TY released from the Chi-TY-AuNP’s was extrapolated using the standard curve of TY at 274 nm (0–100 μg/mL, r^2^ = 0.9934) and plotted the UV-visible absorbance value. Finally, the drug loading efficiency (DLE) was calculated using the following formula:

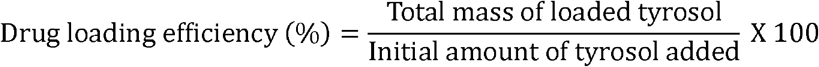

### Strains and Culture Conditions

The strains of *C. albicans* (ATCC SC5314) and *C. glabrata* (MTCC 3019) used in this study were a kind gift from Dr. Navin Kumar, Graphic Era University, Dehradun, India. Yeast extract peptone dextrose agar (YPD) media plates comprising of 2% dextrose, 2% peptone, 2% agar, and 1% yeast extract were used for routine maintenance of strains at 37°C, which were cultured in YPD broth. Rosewell Park Memorial Institute (RPMI) medium was used for *in vitro* biofilm studies.

### Estimation of Minimum Inhibitory and Fungicidal Concentration

Minimum inhibitory concentration (MIC) of Chi-TY-AuNP’s against *C. albicans* and *C. glabrata* planktonic growth was studied in the flat-bottom 96-well multitier plate (MTP) as mentioned in M27-A2 micro broth dilution guidelines of CLSI.^21^ Briefly, 100 µL of cell suspension was obtained from a suspension of log-phase cells (2.5 × 10^3^ cells/mL) prepared using RPMI medium and added into the wells of MTP. 100 µL of RPMI medium containing Chi-TY-AuNP’s in various concentrations 0, 25, 50, 100, 200, 400 and 800 μg/mL was later added into the MTP. Absorbance was taken at 600 nm after 48 h of incubation at 37°C. The inhibition of growth by Chi-TY-AuNP’s was described in MIC_80_, in comparison to control. Chi-TY-AuNP’s inhibited 80% growth of both *Candida* spp. (*C. albicans* and *C. glabrata*). Minimum fungicidal concentration (MFC) of Chi-TY-AuNP’s was determined by spotting 5 µL of MIC sample from MTP on YPD medium plates, followed by incubation at 37°C for 18 1. h. The plates were then photographed for growth analysis. The concentration at which no growth observed was considered as MFC.

### Germ Tube Formation Assay

To study the development of germ tubes in *C. albicans*, protocol earlier described by Gupta et al. was followed.^22^ Briefly, log-phase cells incubated for 4 h at 37°C, with and without Chi-TY-AuNP’s (200 μg/mL) in YPD medium supplemented with 10% FBS. Inhibition of *C. albicans* hyphal development was visualized using a fluorescence microscope (EVOS-FL, Advanced Microscopy Group, USA) at 60X and compared with the positive and negative *C. albicans* control groups.

### Effect of Chi-TY-AuNP’s on *Candida* Biofilm Inhibition and Eradication

The method to determine the effectiveness of Chi-TY-AuNP’s in the inhibition and eradication of *Candida* (*C. albicans* and *C. glabrata*) biofilm was adapted from Gupta et al.^22^ After attaining the log phase, cells of both *Candida* spp. were suspended in PBS (pH 7.0) to attain a cell count of 1×10^7^ cells/mL, following which 100 μL of cell suspension was added to each well of MTP. For the attachment of cells to the MTP wells, the suspension was subjected to incubation at 37°C for 1.5 h. MTP wells were then washed two times with PBS, followed by the addition of RPMI (200 µL) media containing different concentrations of Chi-TY-AuNP’s (0, 25, 50, 100, 200, 400 and 800 μg/mL) for biofilm formation assay. XTT reduction assay was employed to quantify the developed biofilms, followed by 24 h incubation period. For biofilm eradication studies, mature biofilms of both *Candida* spp. were developed in MTP for 48 h. After incubation, media containing Chi-TY-AuNP’s was added and after 24 h of incubation, plates were washed with PBS. Biofilm inhibition and eradication efficacy of Chi-TY-AuNP’s were described in terms of biofilm inhibitory concentrations 80 (BIC_80_) and biofilm eradication concentration 80 (BEC_80_), at which 80% biofilm growth was inhibited.

### Field Emission Scanning Electron Microscopy (FESEM) Analysis

Morphological alterations in Chi-TY-AuNP’s treated *Candida* (*C. albicans* and *C. glabrata*) biofilms were visualized using FESEM. The biofilms were developed on a polystyrene disc of 1 cm^2^, placed in a 24-well plate, incubated with FBS for 24 h. The wells were added with 1 × 10^7^ cells/mL of cell suspension and were incubated at 37°C for 48 h. Later, each well having PBS washed discs, were added with RPMI containing Chi-TY-AuNP’s. Discs were washed with PBS, incubated for 4 h in the dark in 2.5% glutaraldehyde, and then dehydrated using a gradient of ethanol. Samples were air-dried and mounted on stubs for gold sputtering. Visualization of samples was performed using FESEM (voltage-20 kV; magnification 1000 X-5000 X).^22^

### Fluorescence Microscopy Analysis

Live and dead cells in Chi-TY-AuNP’s treated biofilms of both *Candida* spp. were visualized under a fluorescence microscope using fluorescein diacetate (FDA) and propidium iodide (PI) staining. For this study, biofilms of both *Candida* spp. were developed in MTP, as described earlier in the presence of Chi-TY-AuNP’s. The biofilms were washed with PBS after treatment with Chi-TY-AuNP’s (100 μg/mL) and stained with the FDA and PI at a concentration of 2 µg/ mL and 0.6 µg/mL, respectively. After 20 min of incubation in the dark, PBS washed wells were visualized under a fluorescence microscope at 40X magnification.

### Determination of Ergosterol Content

For spectrophotometric determination of ergosterol concentration in the cell membrane, the log phase cells of both *Candida* spp. were incubated in sabouraud dextrose broth consisting of TY, Chi-AuNP’s and Chi-TY-AuNP’s for 20 h at 37°C.^22^ Subsequently, the cells were pelleted down for 5 min at 6000 rpm. The wet weight was measured after washing the cell pellet with Milli-Q water, followed by dissolving in 3 mL of 25% alcoholic potassium hydroxide (lysing agent) and vortexed. Incubation of the cell suspension was done in a water bath for 1 h at 85°C, and a mixture of n-heptane and distilled water (1:3 ratio) was added, followed by vigorous vortexing. The mixture was left undisturbed for 20 min at room temperature. The sterol containing the heptane layer was pipetted gently into the glass tube and was stored at –20°C. After that, a mixture of absolute ethanol (100 μL) and sample (20 μL) was added to analyze the sterol extracts using a UV-visible spectrophotometer (230-300 nm). Ergosterol content was quantified with the help of the following equation:

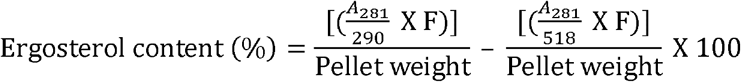

Where the dilution factor, E value for crystalline ergosterol, and E value for 24(28)-dehydro ergosterol were represented by F, 290, and 518, respectively.

### Reactive Oxygen Species (ROS) Generation Assay

The level of ROS in both *Candida* spp. biofilms upon exposure to TY, Chi-AuNP’s and Chi-TY-AuNP’s were determined using 2,7-dichlorodihydrofluoroscein diacetate (DCFDA) and PI.^22^ Briefly, 48 h of mature biofilms were treated with TY, Chi-AuNP’s, and Chi-TY-AuNP’s for 4 h. A mixture of DCFDA (10 μM) and PI (1 mg/mL) was added to the MTP wells with biofilms. After 30 min of incubation in the dark, the fluorescence of DCFDA (λ_ex_ = 520 nm and λ_em_= 485 nm) and PI (λ_ex_ = 617 nm and λ_em_= 543 nm) was measured to assess the level of ROS. Further, a fluorescence microscope was used to capture the microscopic fluorescent images at 40X magnification.

### Estimation of Biochemical Composition of ECM

For biochemical characterization of ECM, *Candida* biofilms were developed in a 24-well plate for 48 h and then treated with TY, Chi-AuNP’s and Chi-TY-AuNP’s for 18 h. After incubation, wells were washed, and the attached biofilms were scrapped with a sterile scrapper in PBS. The biofilms were sonicated in ice using Q700 sonicator (QSonica, 35W), for 5 cycles (30 s each), followed by the centrifugation of suspension for 5 min at 12,000 rpm, after which the supernatant was collected. The estimation of protein and eDNA in ECM was performed with phenol, chloroform, and isoamyl alcohol (PCI) and BCA kit, respectively. For protein estimation, bovine serum albumin was employed as standard, and the absorbance of samples was measured at 562 nm. For eDNA estimation, the samples were added with 1/10^th^ volume of 3 M sodium acetate, followed by the addition of PCI (25:24:1), resulting in the formation of an aqueous layer. The eDNA was precipitated with ethanol (2.5 volumes) and the aqueous layer was collected in a fresh tube. The Nanodrop (Thermo Fisher Scientific, USA) spectrophotometer (A_260/280_) was used to check the purity of eDNA.

### Hydrophobicity Assay

The hydrophobicity of both *Candida* spp. was measured by exposing overnight growing cells to sub-lethal concentration of TY, Chi-AuNP’s and Chi-TY-AuNP’s at 37°C for 24 h. Subsequently, after incubation, the PBS washed cells were suspended in 50 mM sodium phosphate buffer (3 mL; pH 7.0) at 2 × 10^6^ cells/mL concentration after the addition of 500 μL of octane. The cells were then vortexed for 1 min that resulted in the formation of an aqueous layer. The cells of the aqueous layer were observed at OD_600_, and the hydrophobicity index (HI) was calculated using the following formula:

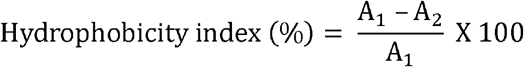

Where A1 and A2 indicate the absorbance of inoculum and aqueous phase, respectively.

### Transcriptional Analysis

The effect of the subinhibitory concentrations of TY, Chi-AuNP’s and Chi-TY-AuNP’s on selected gene expression of *C. glabrata* was assessed with the help of qRT-PCR.^22^ Briefly, log phase *C. glabrata* cells were subjected to 3 h of incubation with TY, Chi-AuNP’s, and Chi-TY-AuNP’s. The RNeasy kit (Qiagen, Germany) was used for extraction of total RNA following the manufacturer’s protocol. Nanodrop spectrophotometer was utilized for qualitative and quantitative analysis of RNA. Verso cDNA synthesis kit (Thermo Fisher Scientific, USA) was used to synthesize cDNA from 1 µg of extracted RNA. The primers of selected genes were procured from Integrated DNA Technologies, India, and RT-PCR was performed using SYBR green master mix for 100 ng of cDNA template and 300 nM of gene-specific primers to make each reaction. For RT-PCR, the following conditions were used: first denaturation cycle was performed at 95°C/3 min, followed by annealing at 60°C/30 s, and extension at 72°C/30 s, the process was repeated for 40 cycles; melting-curve analysis starting from initial temperature 45°C to 95°C, with a gradual increase in 0.5°C/15 s. Melt curve analysis was used as an indicator of primer specificity. The cycle threshold (CT) values of the housekeeping *ACT1* gene were used to normalize the CT values of target genes. ΔΔCT method using 2-ΔΔCT formula was utilized to evaluate the relative expression fold changes.

### Statistical Analysis

All experiments were performed in triplicates, and the values presented as the mean ± standard deviation (SD), obtained from three different observations for each assay. Student’s t-test was used for the statistical analysis, and a value of \**P* < 0.05 was considered statistically significant \*\**P* < 0.01 as highly significant \*\*\**P* < 0.001 as extremely significant.

## RESULTS

### Physicochemical Characterization of Chi-TY-AuNP’s

The physicochemical characterization of Chi-TY-AuNP’s was carried out by analyzing the size, zeta potential, aggregation behavior and chemical interactions were studied with electron microscopy, zetasizer and FTIR analysis, respectively. Synthesis of Chi-TY-AuNP’s was carried out using chitosan, which displays both the stabilizing and reducing properties, and the size of the average nanoparticles was determined, as shown in **Figure 1**. Confirmation of synthesized Chi-AuNP’s was ascertained by UV-visible spectroscopy. The bio-reduction of Au^+^ ions into Au^0^ by chitosan was determined by the change in color from transparent to wine red/ ruby red color (**Figure 1A**). The reaction mixture was allowed to cool at 25°C and analyzed by UV-Vis spectroscopy at subsequent time intervals from 0 h to 168 h at 531 nm. The appearance of ruby red color and consistent sharp peaks at 531 nm confirms the formation of Chi-AuNP’s. The presence of sharp peaks in the visible range is attributed to the excitation of its SPR, which in turn is dependent on the shape and size of the nanoparticles.^19, 20^ Furthermore, no significant variation was observed upon drug loading in the absorption spectra of Chi-TY-AuNP’s (**Figure 1A**). Zeta potential is a significant physiochemical attribute for nanosystems, which plays a crucial role in the drug delivery mechanism (**Figure 1B**). The nanoparticulate system’s solubility, cellular absorption and release rate are also influenced by zeta potential. Our findings showed that Chi-AuNP’s and Chi-TY-AuNP’s possess a charge of +62 mV (**Figure 1D**) and +45.5 mV (**Figure 1D**), respectively (**Figure 1B**).^20^ DLS has been used to estimate the diameter of Chi-TY-AuNPs, and the average hydrodynamic diameter was found 46.96 nm (**Figure 1C**), and low PDI of 0.171 substantiates greater colloidal stability in the aqueous environment.

**Figure 1.**
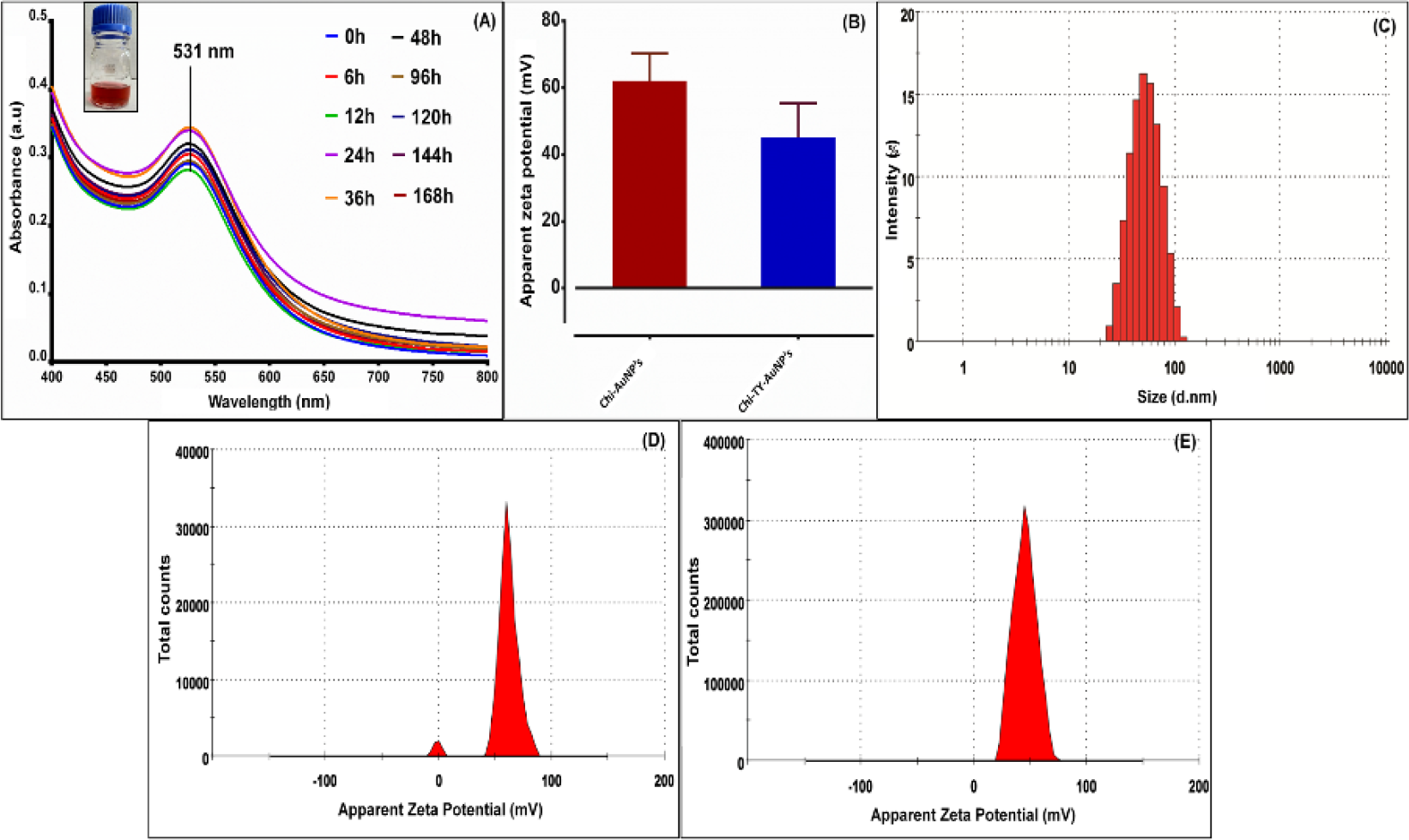
Physicochemical analysis of Chi-TY-AuNP’s (A) UV-visible spectroscopic analysis (B) zeta potential (C) dynamic light scattering (D) apparent zeta potential of Chi-AuNP’s (E) apparent zeta potential of Chi-TY-AuNP’s.

Chi-TY-AuNP’s HRTEM images showed that the nanoparticles were in the range of 10.345±2.684 nm in diameter with a sphere-shaped morphology (**Figure 2A, B, C, D**). AFM investigation revealed the spherical shape of Chi-TY-AuNP’s with an average diameter in the range of 10-15 nm (as shown through nanoparticles color scripting and 3D image) was observed (**Figure 2E, F**). Therefore, from the findings of TEM, AFM, and zeta-potential, we can infer that the size obtained is efficacious in harnessing the antifungal property of Chi-TY-AuNP’s owing to its enhanced permeability and retention (EPR) effect.^20^ The selected area electron diffraction (SAED) pattern of Chi-TY-AuNP’s reveals a transition state of polycrystalline nature. The broad spheres present in Chi-TY-AuNP’s are attributes of the chitosan matrix and rings are made of crystalline gold nanoparticulate (**Figure 2H**).

**Figure 2.**
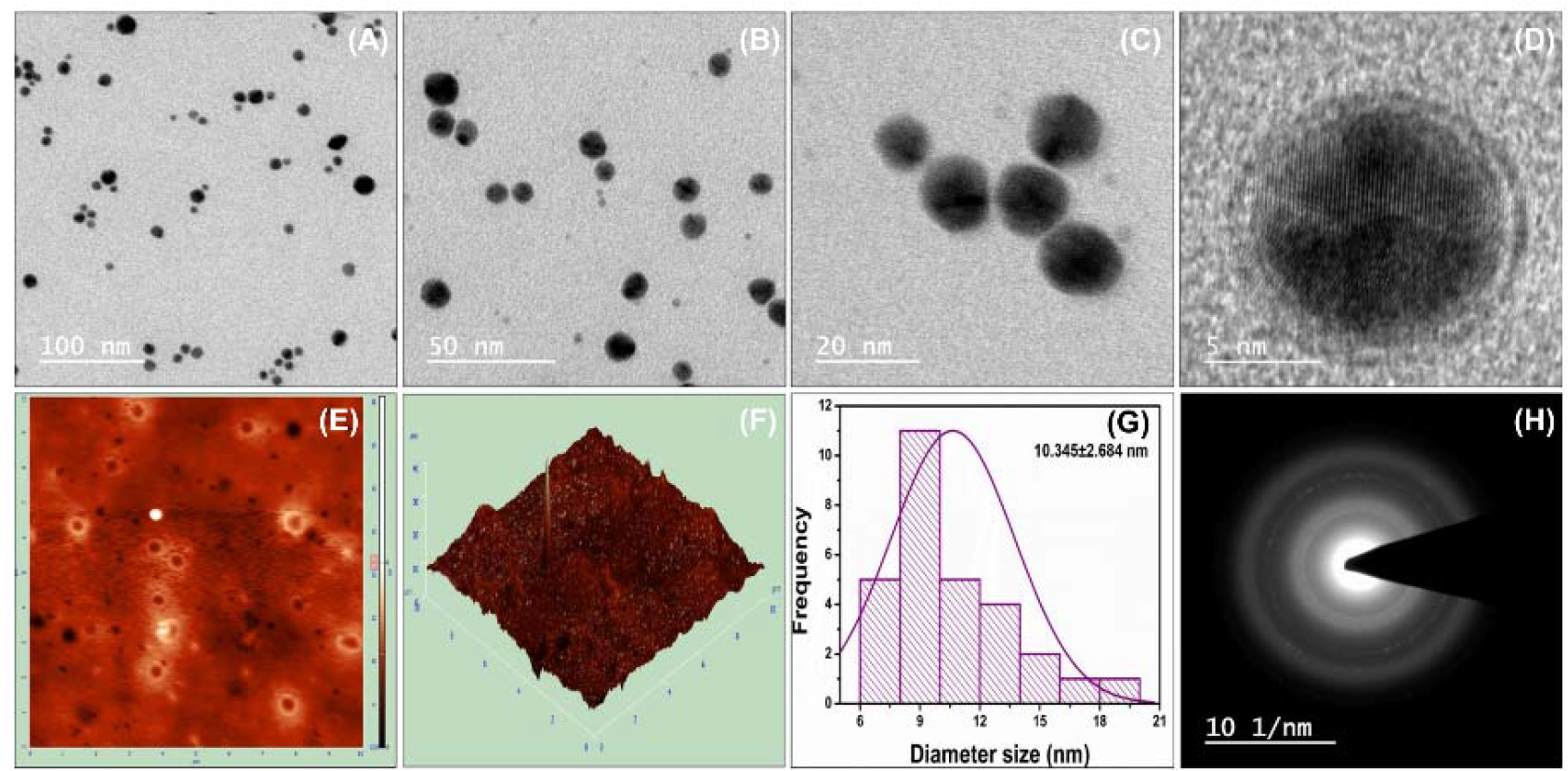
Morphological analysis of Chi-TY-AuNP’s, TEM images at (A) 100 nm (B) 50 nm (C) 20 nm (D) 5 nm (E) 2D AFM (F) 3D AFM (G) particles size distribution curve (H) SAED pattern of synthesized Chi-TY-AuNP’s.

### Fourier-Transform Infrared (FTIR) Analysis

A comparative FTIR analysis was performed for the chemical characterization and functional group validation in Chi-TY-AuNP’s with respect to chitosan and TY (**Figure 3**). During the FTIR analysis, the carbonyl, C–O–NHR, NH_2_ and ammonium, NH_3_ band, OH, and CH deformation in the region of 100-2400 cm^-1^ have been considered as critical analytical peaks. Functional groups assigned to chitosan such as N-H, O-H, and NH_2_ peaks at 3357-3290 cm^-1^ followed with a tiny peak of 2879 assigned to -CH_2_ and -CH_3_ cm^-1^ (**Figure 3A**).^23^ However, the amide II bands (C–N stretching coupled to NH bending) and amide I peak (C=O stretching), as represented in **Figure 3A**, were also observed around 1513 cm^-1^ and 1643 cm^-1,^ respectively.^24^

**Figure 3.**
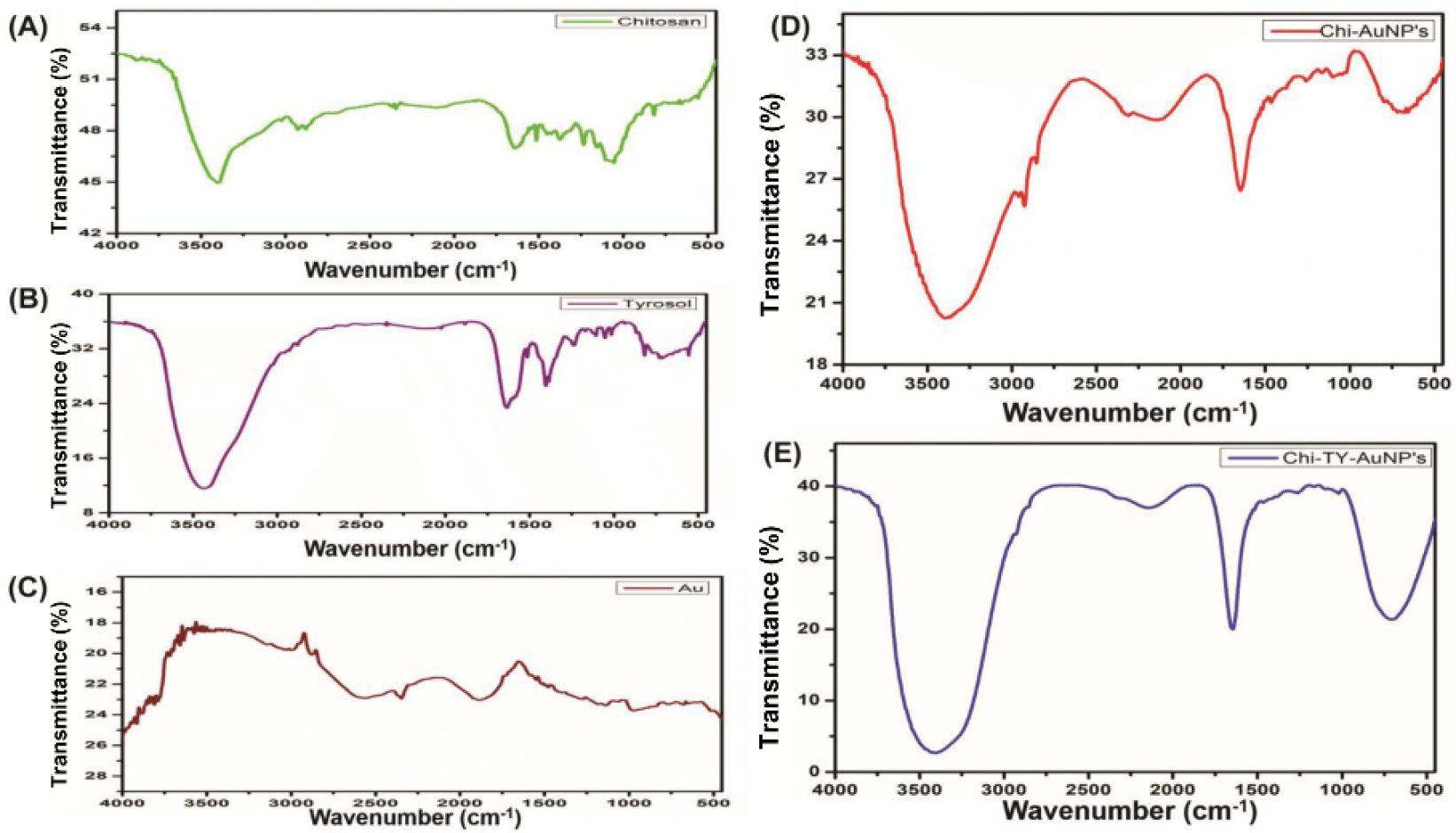
FTIR analysis (A) chitosan (B) tyrosol (C) gold chloride (D) Chi-AuNP’s (E) Chi-TY-AuNP’s.

Nevertheless, the FTIR analysis of TY shows a small peak of C=C aromatic stretching at 1513 cm^-1^. The standard TY peaks are as follows: two bands from 1600-1400 cm^−1^ correlated to C=C (benzene ring), and aromatic C-H (in para position) forms the bending vibration in 900-800 cm^−1^ range (**Figure 3B**).^25^ To ascertain the interactions between chitosan functional groups and gold nanoparticles, the FTIR spectra of Chi-AuNP’s were also assessed. The chemical interactions indicate the shift in the characteristic peaks of the spectrum. The FTIR analysis of Chi-AuNP’s, obtained with nearly identical peaks of chitosan, showing a uniform chitosan deposition over gold nanoparticles (**Figure 3D**).^20^ However, in the Chi-TY-AuNP’s, as in **Figure 3E**, both the peaks of chitosan and TY were evident. This shows the successful adsorption of TY over the surface of Chi-AuNP’s. The apparent disappearance of TY associated peaks substantiates our finding in drug loading efficiency and pH triggered release profile.

### Drug Loading Efficiency

The drug loading efficiency of Chi-TY-AuNP’s was found to be 46.08% as obtained by UV-visible analysis in 100 mg of Chi-TY AuNP’s.

### Chi-TY-AuNP’s Showed Fungicidal Activity Against *Candida* spp

The planktonic growth of both the *Candida* spp. employed in this study was inhibited in a concentration-dependent manner (**Figure 4A**). The MIC_80_ value of Chi-TY-AuNP’s was 200 and 400 μg/mL for *C. albicans* and *C. glabrata* growth, respectively. Chi-TY-AuNP’s effectiveness was substantially greater against the growth of *C. albicans* as compared to *C. glabrata.* Nevertheless, the MFC value of Chi-TY-AuNP’s for both *C. albicans* and *C. glabrata* was 800 μg/mL (**Figure 4B**).

**Figure 4.**
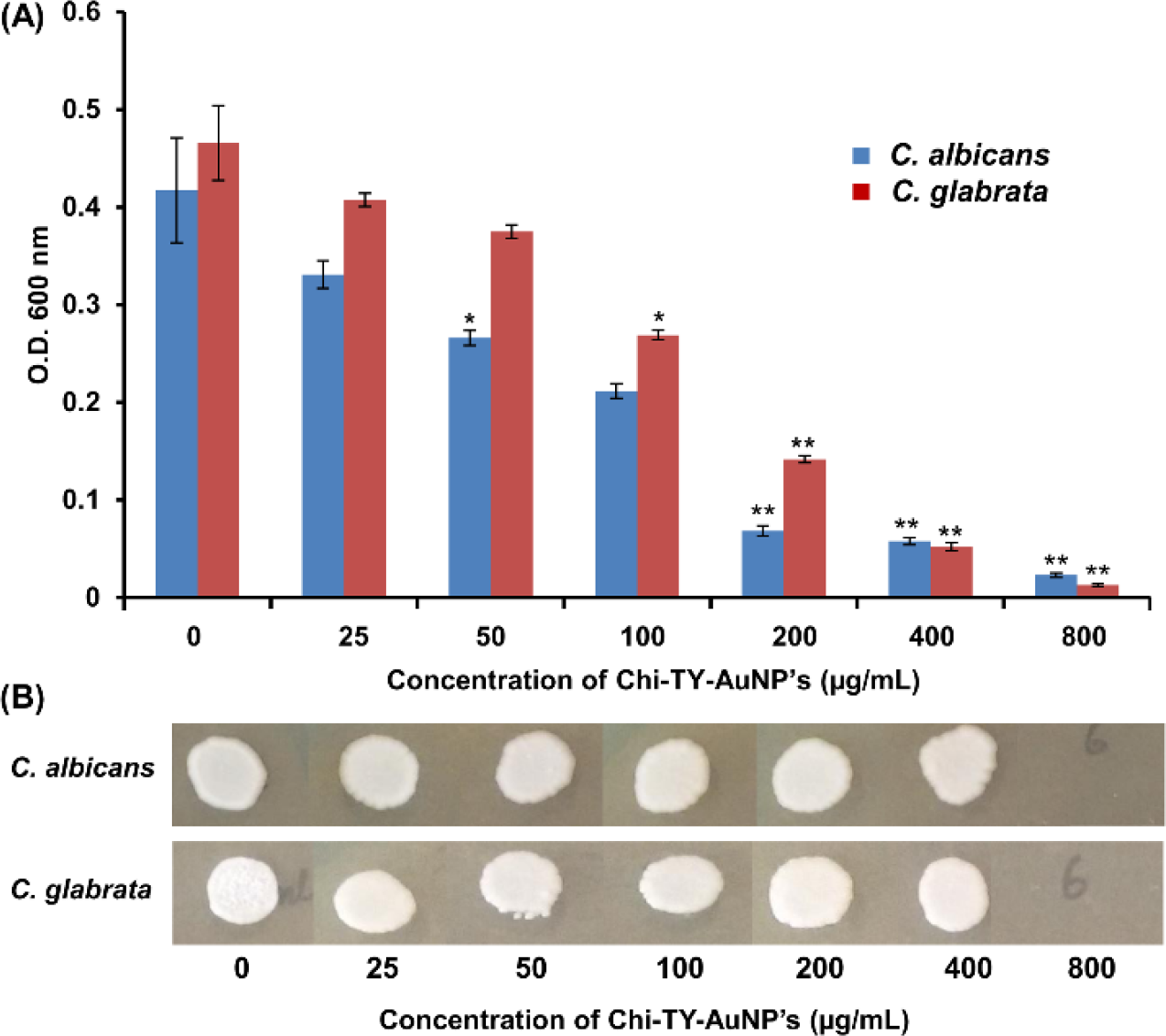
Determination of (A) MIC and (B) MFC of Chi-TY-AuNP’s against *C. albicans* and *C. glabrata* planktonic cells, error bars represent SD (n = 3), \**P* < 0.05 considered as statistically significant.

### Effect of Chi-TY-AuNP’s on *Candida* Biofilm

The concentration-dependent activity of Chi-TY-AuNP’s enabled it to inhibit biofilm development and eradicate the mature biofilms of *C. albicans* and *C. glabrata* (**Figure 5**). The BIC_80_ of Chi-TY-AuNP’s against both *Candida* spp. viz., *C. albicans* and *C. glabrata* was 200 and 400 μg/mL, respectively. At the highest concentration (800 μg/mL), Chi-TY-AuNP’s inhibited 95.98% and 96.34% of *C. albicans* and *C. glabrata* biofilms, respectively (**Figure 5A**). In contrast, Chi-AuNP’s showed 11.36% inhibition of *C. albicans* biofilm and 14.21% inhibition of *C. glabrata* biofilm (Data not showed). The biofilm eradicating efficiency of Chi-TY-AuNP’s, measured in terms of BEC_80_, was 400 μg/mL against both the *Candida* spp. used in this study (**Figure 5B**). The biofilm eradicating efficacy of Chi-TY-AuNP’s was found to be equal against both *C. albicans* and *C. glabrata* biofilms, and the BEC_80_ value was found to be 800 μg/mL for both spp. Further, the morphological alterations in Chi-TY-AuNP’s treated *C. albicans* and *C. glabrata* biofilm cells were visualized by FESEM analysis (**Figure 5C**). The FESEM micrographs of untreated *Candida* biofilm showed a compact network of *C. albicans* hyphal cells, whereas *C. glabrata* cells appeared as healthy elongated and oval with no alteration in the surface topology of cells (**Figure 5C, micrograph i, iii**). However, the biofilm of *C. albicans*, in the presence of 400 μg/mL Chi-TY-AuNP’s, exhibited the absence of hyphal network and wrinkled cells while *C. glabrata* treated biofilm showed cells with pores on the surface (**Figure 5C, micrograph ii, iv**).

**Figure 5.**
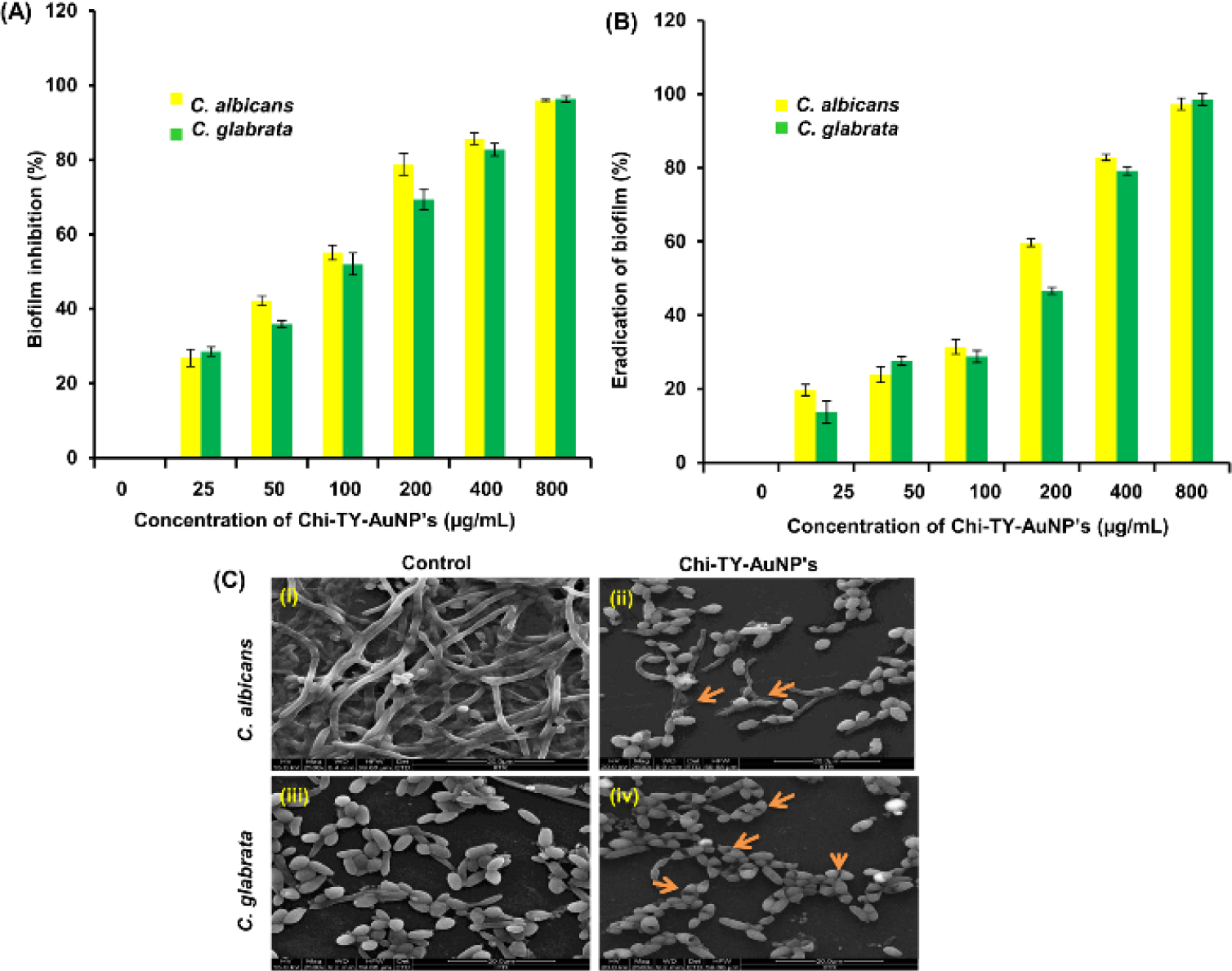
Effect of Chi-TY-AuNP’s on *C. albicans* and *C. glabrata* (A) biofilm eradication and (B) biofilm inhibition (C) FESEM images representing the Chi-TY-AuNP’s induced morphological changes in *C. albicans* and *C. glabrata*, (i) and (iii) Untreated biofilms of *C. albicans* and *C. glabrata* respectively (ii) and (iv) Chi-TY-AuNP’s treated biofilms of *C. albicans* and *C. glabrata* respectively, error bars in graph represent SD (n = 3), magnification 1000-5000X.

### Chi-TY-AuNP’s Inhibited *C. albicans* Germ Tube Formation

Since, Chi-TY-AuNP’s worked extremely well against *Candida* biofilms, to explore the possible antibiofilm mode of action of synthesized nanoparticles and their impact on hyphae was also investigated. But, this study was limited to *C. albicans* because *C. glabrata* do not form germ tubes. **Figure S1** clearly showed the inhibition of germ tube development in *C. albicans* by Chi-TY-AuNP’s at subinhibitory concentration (200 μ L). The germ tubes were induced by exposing cells to 10% FBS. Negative control cells remained in yeast and budding yeast form, while positive control showed long hyphae formation (**Figure S1A, B**). At subinhibitory concentration, Chi-TY-AuNP’s completely inhibited yeast to hyphae transformation and reduced the number of cells (**Figure S1C**). Chi-AuNP’s did not show any alteration in *C. albicans* morphology (**Figure S1D**).

Besides hyphal development, surface hydrophobicity also has a role in biofilm establishment and hence, the effect of Chi-TY-AuNP’s in modulating hydrophobicity of the cells was evaluated in terms of HI using a two-phase system. The hydrophobicity evaluation results suggested no significant change in the HI value of both the *Candida* spp. compared to control cells upon Chi-TY-AuNP’s treatment. While the HI value of *C. albicans* cells increased in response to TY treatment, the value remains nearly the same in Chi-AuNP’s and Chi-TY-AuNP’s treated cells (**Figure S2**). Therefore, hydrophobicity was not observed to be responsible for mediating the antibiofilm activity of Chi-TY-AuNP’s.

### Effect of Chi-TY-AuNP’s on Viability of *Candida* Biofilm Cells

Further, FDA-PI staining was used to assess the live and dead cells in Chi-TY-AuNP’s treated *C. albicans* and *C. glabrata* biofilms and visualized using a fluorescence microscope (**Figure 6**). The FDA binds with the cell membrane polysaccharides of living cells and emits green fluorescence while lysed/ dead cells fluoresce red due to PI’s binding with DNA. *C. albicans* control biofilm and treated with Chi-AuNP’s emitted green fluorescence indicating no cell death, whereas Chi-TY-AuNP’s treated cells exhibited both red as well as green fluorescence indicating cell lysis (**Figure 6A**). Likewise, no red fluorescence was observed in *C. glabrata* control biofilm and biofilm treated with Chi-AuNP’s; however, intense red fluorescence with dull green light was observed in Chi-TY-AuNP’s treated biofilm (**Figure 6B**).

**Figure 6.**
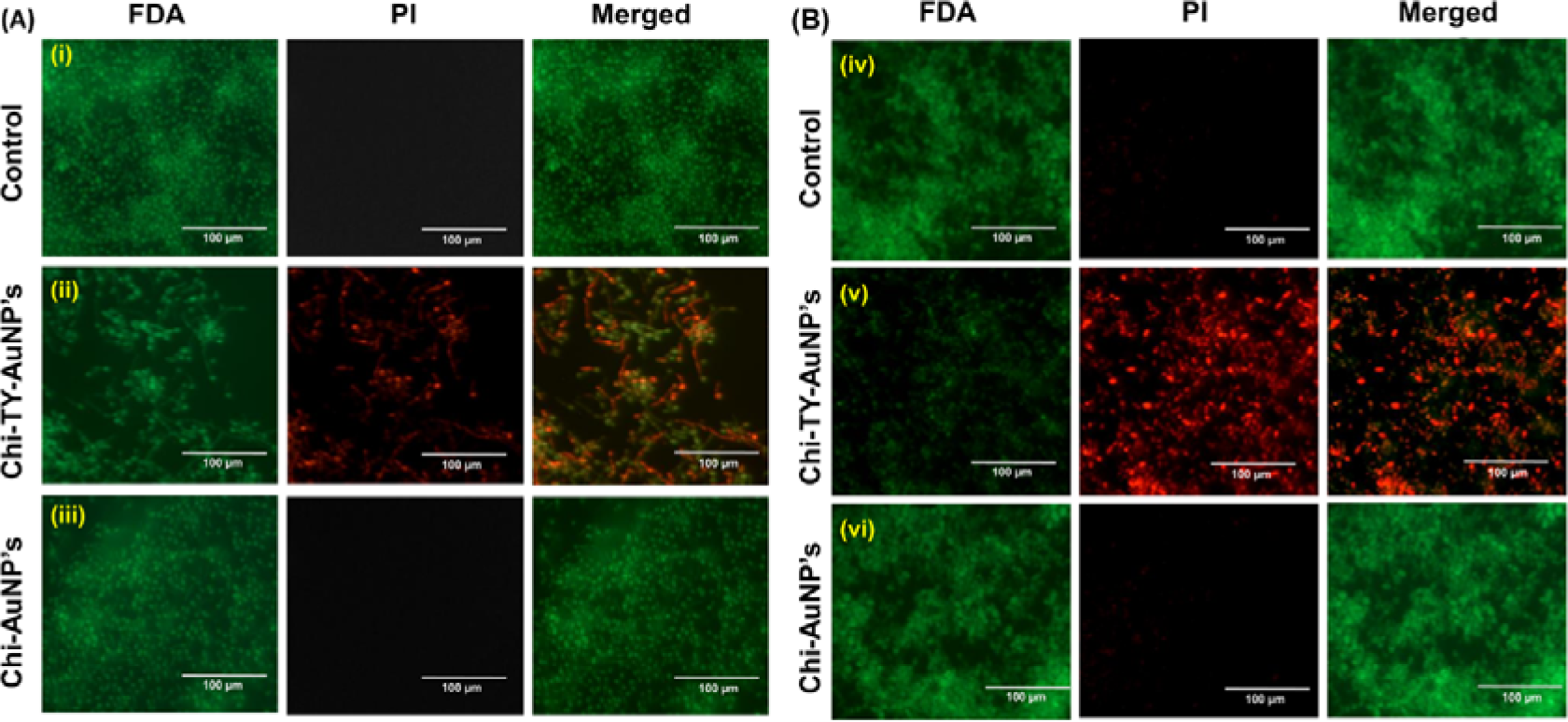
FDA/PI stained live and dead cells (A) *C. albicans* (B) *C. glabrata* (i) and (iv) control (untreated) *C. albicans* and *C. glabrata* cells respectively (ii) and (v) Chi-TY-AuNP’s treated *C. albicans* and *C. glabrata* cells respectively (iii) and (vi) Chi-AuNP’s treated *C. albicans* and *C. glabrata* cells respectively, magnification: 40X, scale bar: 100 μm.

### Effect of Chi-TY-AuNP’s on Biofilm Extracellular Matrix (ECM)

To gain insight into Chi-TY-AuNP’s mediated effects on biochemical components of *C. albicans* and *C. glabrata* biofilm ECM, the spectrophotometric study was performed. Insignificant changes were observed in the protein content of *C. albicans* as compared to control in Chi-AuNP’s and Chi-TY-AuNP’s while significantly increased in TY. However, in *C. glabrata*, the protein content was considerably reduced in Chi-TY-AuNP’s as compared to control and remained unchanged in the rest of the samples (**Figure 7A**). The eDNA content of *C. glabrata* ECM was relatively much higher than *C. albicans* control biofilm. The eDNA content in *C. albicans* biofilm was increased in TY, whereas substantially decreased in Chi-TY-AuNP’s, and remained unchanged in Chi-AuNP’s. In *C. glabrata*, the content of eDNA was decreased in all samples with the highest reduction in Chi-TY-AuNP’s (**Figure 7B**).

**Figure 7.**
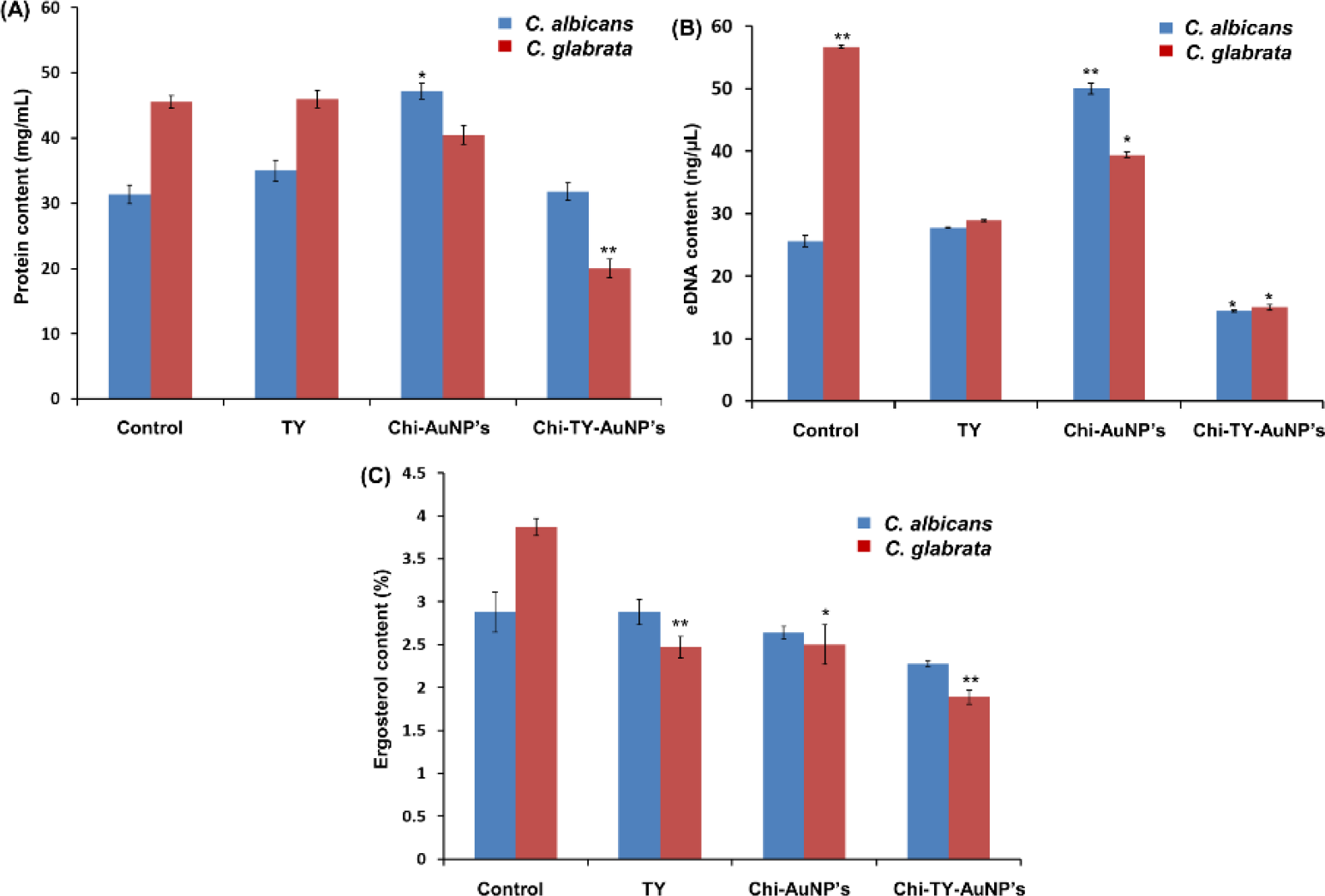
Biochemical quantification of TY, Chi-AuNP’s and Chi-TY-AuNP’s treated *C. albicans* and *C. glabrata* (A) estimation of protein (B) estimation of eDNA (C) estimation of ergosterol, error bars represent SD (n = 3), \**P* < 0.05 considered as statistically significant.

### Analysis of ROS Production in *Candida* Biofilms Treated With Chi-TY-AuNP’s

ROS generation is one of the most widely adopted strategies of drug molecules to mount antimicrobial activity and sometimes promote cell death. To estimate the amount of ROS produced by TY, Chi-AuNP’s and Chi-TY-AuNP’s in *Candida* biofilms, PI and DCFDA were used. PI binds with DNA manifesting cell lysis while DCFDA determines the ROS level generated within the cells. *C. albicans* biofilm treated with Chi-TY-AuNP’s and Chi-AuNP’s exhibited significant elevated level of ROS in comparison to control; however, ROS elevation was more pronounced in *C. glabrata* biofilm (**Figure 8A, B**). The interaction of PI with DNA of lysed cells determined the detrimental effect of ROS accretion on cells; high fluorescence intensity of intercalated PI was observed in both the *Candida* spp. biofilms treated with Chi-TY-AuNP’s (**Figure 8C, D**).

**Figure 8.**
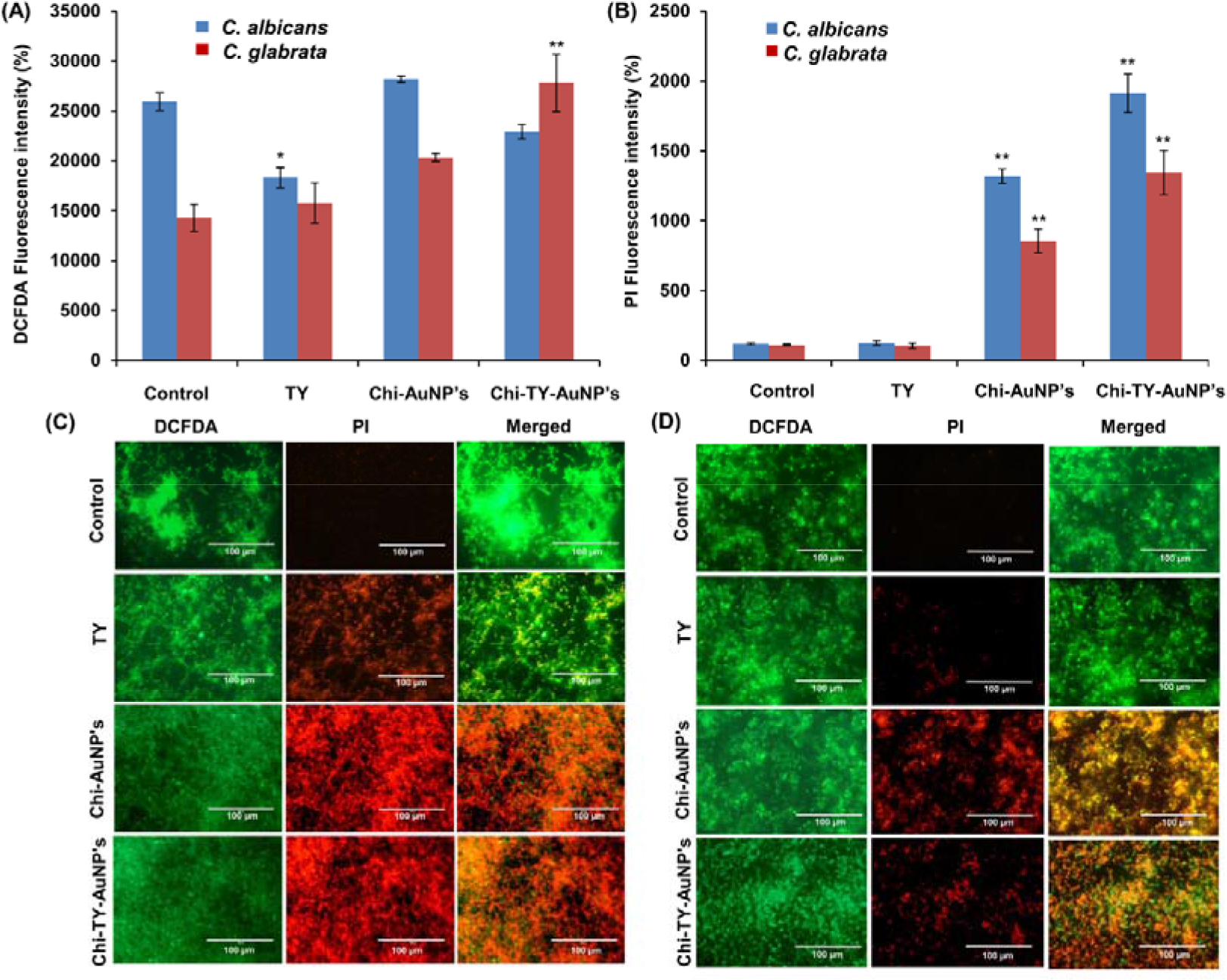
Measurement of ROS generation in C. *albicans* and *C. glabrata* cells exposed to TY, Chi-AuNP’s, and Chi-TY-AuNP’s. ROS level is represented in terms of fluorescence intensity of (A) DCFDA and (B) PI; and microscopic fluorescence images of (C) *C. albicans* and (D) *C. glabrata*, error bars in graph represent SD (n = 3), \**P* < 0.05 considered as statistically significant; magnification: 40X, scale bar: 100 μm.

### Chi-TY-AuNP’s Differentially Modulated *C. glabrata* Transcriptional Expression

The cell membrane of the *Candida* is the first line of defense against any drug molecule, like all other organisms. Ergosterol is a critical component of the *Candida* cell membrane, which plays a remarkable role in its susceptibility to drug molecules and, thus, a primary target for antifungal drugs. Therefore, the change in ergosterol content of the plasma membrane was estimated upon exposure of both the *Candida* spp. to Chi-TY-AuNP’s. *C. albicans* exhibited no change in ergosterol content in the presence of Chi-TY-AuNP’s, while *C. glabrata* showed decreased ergosterol concentration in comparison to control (**Figure 7C**).

Therefore, the transcriptional expression study of the ergosterol pathway and ABC transporters was performed in *C. glabrata* cells. The transcriptional expression of sterol importer (*AUS1*), multidrug transporter (*CDR1*), 1,3-β-glucan synthase (*FKS1*), ergosterol synthetic pathway (*ERG11, ERG3, ERG2, ERG10*, and *ERG4*) and GPI-anchored cell wall protein (*KRE1*) genes was studied using RT-PCR. Except for *ERG11*, expression of *AUS1, KRE1*, *FKS1,* and all genes of the ergosterol pathway were significantly downregulated upon Chi-TY-AuNP’s treatment. (**Table 1**). The trend in the gene expression pattern of TY treated *C. glabrata* cells was similar to Chi-TY-AuNP’s treated cells. However, the fold change value of TY treated *C. glabrata* cells was lesser than the Chi-TY-AuNP’s treated cells.

**Table 1.** Up and downregulation of *C. glabrata* genes in response to the subinhibitory concentration of Chi-TY-AuNP’s.

| *C. glabrata*<br>*Genes* | Function | Fold expression |  |  |
| --- | --- | --- | --- | --- |
|  |  | TY | Chi-AuNP's | Chi-TY-AuNP's |
| *CDR1* | ABC multidrug-transporter regulated by Pdr1p | 1.19 | 12.15 | 10.00 |
| *ERG2* | C-8 sterol isomerase activity | No | 1.4 | -2.0 |

|  |  | change |  |  |
| --- | --- | --- | --- | --- |
| *ERG3* | C-5 sterol desaturase activity | -1.67 | 1.1 | -2.0 |
| *ERG4* | C-24 sterol reductase activity | 1.7 | 2.1 | -2.0 |
| *ERG10* | Acetyl-CoA acetyltransferase have a role in sterol biosynthesis | -1.42 | 2.00 | -3.34 |
| *ERG11* | Cytochrome P-450 lanosterol 14-alpha-demethylase role in ergosterol synthesis | 1.2 | 43.41 | 12.2 |
| *AUS1* | ABC transporter involved in sterol uptake | -2.0 | -1.42 | -10.0 |
| *KRE1* | Role in cell wall biogenesis and organization | -2.5 | -2.0 | -2.5 |
| *FKS1* | Component of 1,3-beta-glucan synthase | -2.5 | -3.34 | -10 |

Genes showing a fold expression $\geq 1.5$ were only considered to assess the changes. Fold expression of 1.5-5.0 was considered as significant.

## DISCUSSION

Members of genus *Candida* are commensal, opportunistic fungal pathogens with biofilm-forming ability as the most prominent virulence feature responsible for the evolution of MDR *Candida* strains and therapeutic failure of conventional antifungals.^11^ Due to its recalcitrant feature, complete removal of biofilm is a grave challenge. Nanocarrier drug delivery systems emerged as a promising strategy due to their biofilm barrier penetrating capacity owing to their smaller size and their localization into cellular and subcellular compartments for site-specific antibiotic delivery. Engineering of biopolymer catalyzed metal nanoparticles with antibiotics/ small molecules will improve their intracellular delivery and antibiofilm effect via preventing microorganism’s surface adherence and internalization into microbial cells resulting in the destruction of intracellular architecture.^26, 27^ Polymeric metal nanoparticles offer excellent drug loading efficiency manifesting greater therapeutic index and improved pharmacokinetic profile in low dose compared to the free form of the drug with minimal associated side-effects.^28^

TY, as an indigenous QSM, is well known to induce the yeast-to-hyphae transformation via concentration-dependent fungal growth in a culture medium. Recent investigations showed the fungicidal and antibiofilm effects of exogenously administered TY in different *Candida* spp., but these studies were insufficient to prove the therapeutic potential of TY in complete eradication of biofilm and its mechanistic insight.^7,29^ In view of the antimicrobial property and biocompatibility of both chitosan and gold combined with the increased susceptibility of TY towards *Candida,* we synthesized Chi-TY-AuNP’s and evaluated its antifungal and antibiofilm activities against *C. albicans* and *C. glabrata*. Further, we unraveled the mechanistic insights of Chi-TY-AuNP’s by assessing their effect on ROS generation, cell surface hydrophobicity, ECM composition and membrane ergosterol content in biofilms of both the *Candida* spp. along with this, transcriptional expression of selected *C. glabrata* genes was also evaluated.

The physicochemical analysis revealed the spherical architecture of synthesized Chi-TY-AuNP’s in size range of 10.345 ± 2.684 nm in diameter (**Figure 2G**). The higher the zeta potential, the greater will be the stability of colloidal nanoparticles due to electrostatic repulsion.^13^ In our study, the zeta potential of Chi-TY-AuNP’s was found significantly higher attributed to the cationic characteristic of chitosan, signifying greater colloidal stability (**Figure 1B**). Conclusively from the findings of zeta-potential, HRTEM and AFM we can infer that the high positive surface charge and smaller size obtained is efficacious in harnessing the antifungal property of Chi-TY-AuNP’s owing to its EPR effect towards the negatively charged cytoplasmic membrane.^20^ Further, FTIR analysis showed the functionalization of TY over the surface of nanoparticles (**Figure 3E**), and high surface to volume ratio of Chi-TY-AuNP’s signifying high drug payload.

Chi-TY-AuNP’s has efficiently reduced the fungal growth and killed the sessile as well as planktonic cells of both *Candida* spp. in a concentration-dependent manner (**Figure 4A, B**). The observed fungicidal action of Chi-TY-AuNP’s may be a result of their smaller size and the ability of gold to its preferential binding with the cell surface, leading to intracellular localization of Chi-TY-AuNP’s within the cytoplasm (**Figure 9A**). Furthermore, the fungicidal effect of Chi-TY-AuNP’s may be attributed to the interaction of –NH_2_ groups present on the polycationic chitosan with the negatively charged cytoplasmic membrane via electrostatic bond formation. This interaction leads to the disruption of the fungal cell membrane resulting in leakage of intracellular components.^30^ The higher positive surface charge of Chi-TY-AuNP leads to strong electrostatic interaction between nanoparticles and fungal cells, resulting in increased nanoparticle penetration inside the microbial cell, corroborating higher fungicidal activity.^13, 31^ Besides, the synergistic antimicrobial effect of chitosan and gold altered the intercellular signaling pathways and an imbalance of cell metabolism leading to loss of fungal cell viability. These results are in line with the previous studies where TY alone or with other antifungals upon exogenous administration have inhibited the biofilm formation in different *Candida* spp., including *C. albicans* and *C. glabrata*.^7, 29, 32^

**Figure 9.**
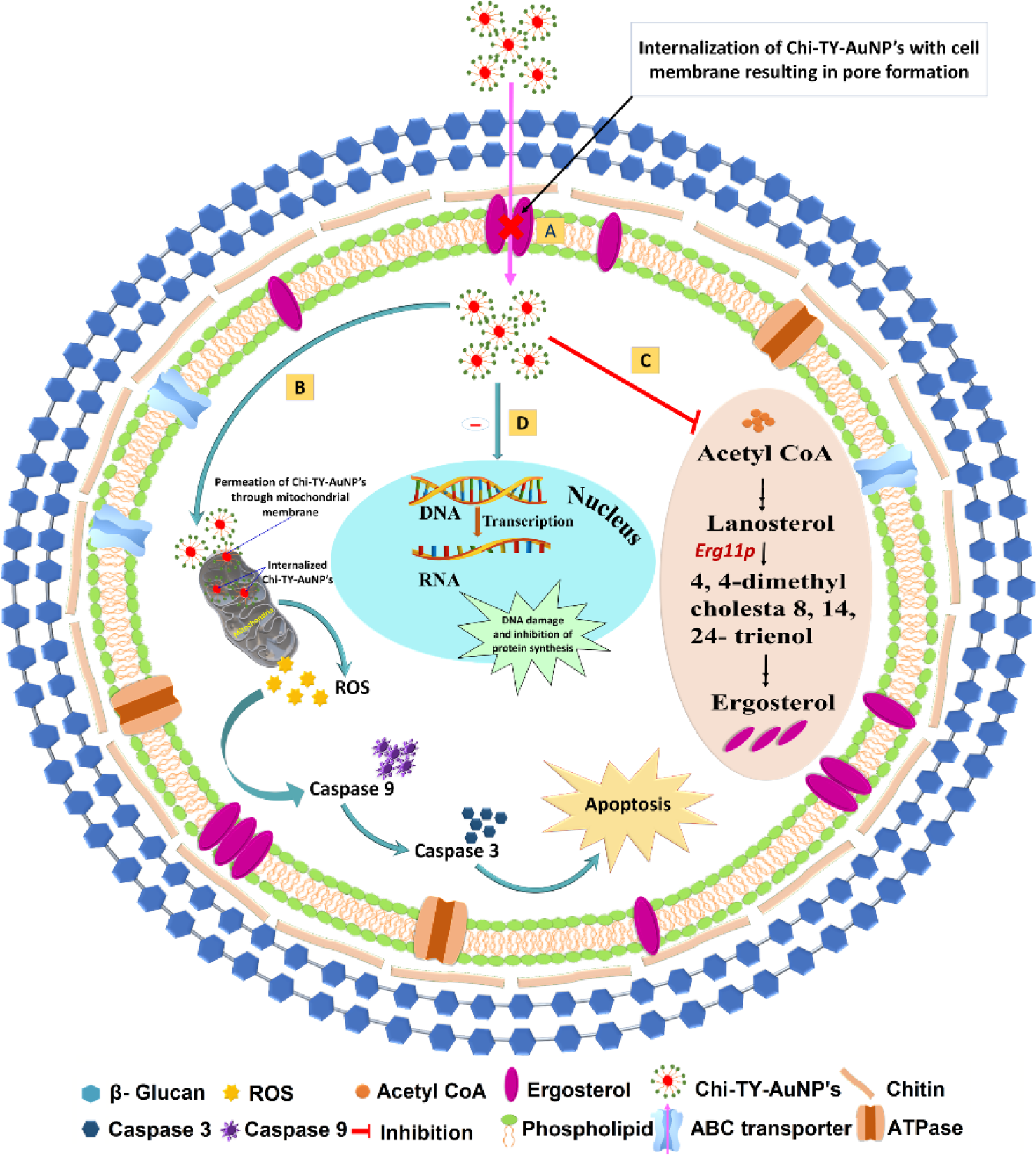
Schematic representation of possible antifungal and biofilm inhibition mechanism of Chi-TY-AuNP’s. (A) cellular uptake and internalization of Chi-TY-AuNP’s with cell membrane resulting in pore formation, (B) Chi-TY-AuNP’s mediated ROS generation leading to cell apoptosis, (C) inhibition of ergosterol and glucan biosynthesis resulting in increased susceptibility of the cell membrane to external stress, (D) modulation of ECM composition leading to inhibition of cellular communication between biofilm cells and finally resulting in architectural collapse.

Chi-TY-AuNP’s efficiently inhibited and eradicated the biofilms of *C. albicans* and *C. glabrata* in a concentration-dependent manner. (**Figure 5A, B**). In the present study, the Chi-TY-AuNP’s concentration, which inhibited the biofilm formation of both the *Candida* spp. was much lower than that of TY used in earlier studies.^7, 29^ Strong electrostatic attractive forces and high positive charge exhibited by the –NH_2_ groups of polycationic chitosan present in Chi-TY-AuNP’s are expected to interact with the negatively charged cell membrane leading to the migration and internalization of Chi-TY-AuNP’s to the subcellular environment of cytoplasm.^33^ This cell surface-nanoparticles interaction is not only mediated by the physicochemical characteristics of nanoparticles such as size, surface to volume ratio, surface functionalization and zeta potential etc. the structural components of biofilm matrix also equally contributed to this event.^34^ Furthermore, internalized Chi-TY-AuNP’s synergistically interacted with the cellular components (DNA, RNA and proteins), resulting in the inhibition of protein synthesis and cell respiration. All these events might be collectively responsible for the inhibition of early formation of biofilms and eradication of mature biofilms.^33^ Considering the potency of Chi-TY-AuNP’s for complete eradication of biofilm of both the *Candida* spp. which may be the net therapeutic effect of tyrosol, chitosan and gold coupled in a nanocarrier system. The morphological alterations and germ tube inhibition were also observed in response to Chi-TY-AuNP’s treated *C. albicans* and *C. glabrata* cells (**Figure S1, 5C**). The FESEM observations are in line with the previous study conducted by Arias et al. where similar morphological distortions in TY (50-200 mmol^-1^ L^-1^) exposed *C. albicans* biofilms were evident.^29^ The assessment of cell damage and topological distortions mediated by Chi-TY-AuNP’s on treated biofilms were carried out by FDA-PI staining. All these observations supported the fungicidal and antibiofilm efficacy of Chi-TY-AuNP’s against *Candida*.

Adhesion of microbes to the cell surface is an early stage event and a major determinant in biofilm formation, which depends on several physiological factors, including physical and chemical surface characteristics of the cell, nutritional growth factors and viability of microbial cells. Early inhibition of microbial adhesion has been shown to prevent further formation of biofilms.^35^ Cell surface hydrophobicity has a significant role in the interactions between bacterium and host cell and further in microbial biofilm matrix formation. Targeting HI is a novel strategy to combat biofilm formation and maturation.^36^ In our study, we did not observe any significant change in the HI value of both *Candida* spp. upon Chi-TY-AuNP’s treatment compared to control. (**Figure S2**) suggesting Chi-TY-AuNP’s is not participating in any kind of cell surface hydrophobicity related activity in *Candida* cells.

The antimicrobial resistance of *Candida* biofilms is multifactorial, with additional protection facilitated by the biofilm ECM assembly. Structural components of *Candida* ECM such as hydrolytic enzymes, polysaccharides, protein, β-glucans and eDNA contribute to tissue penetration and invasion and maintenance of structural integrity and stability of *Candida* biofilms.^37^ These structural components of ECM collectively restrict the diffusion of antifungals to biofilms leading to reduced therapeutic effectiveness and incomplete eradication of biofilms.^38^ Since ECM components are major contributing factors in the formation of *Candida* biofilms, we studied the effects of Chi-TY-AuNP’s on *C. albicans* and *C*. *glabrata* biofilms ECM to determine their interference with ECM components. We observed a significantly reduced protein content in Chi-TY-AuNP’s treated *C. glabrata* biofilm, while this effect was insignificant in *C. albicans* (**Figure 7A**). The functional role of eDNA has been correlated with the hyphal growth in *C. glabrata,* while it protects and stabilizes mature biofilms of *C. albicans*.^37, 39^ Additionally, eDNA acts as a master regulator of biofilm formation and antifungal resistance.^37^ In our findings, the substantially decreased eDNA content of both the Chi-TY-AuNP’s treated *Candida* spp. biofilms (**Figure 7B**) can be correlated with the reduced hyphal development as seen in the germ tube formation assay and biofilm susceptibility to Chi-TY-AuNP’s (**Figure S1**). eDNA being polyanionic, its electrostatic interaction with polycationic chitosan present in Chi-TY-AuNP’s alongside hydrophobic interactions of leached gold covalently bound to Chi-TY-AuNP’s with the oxygen and nitrogen atoms of eDNA results in eDNA damage and inhibition of biofilm formation.^40, 41^ Another possible explanation is that gold has the affinity for the proteins and phosphate groups present in the DNA, and their interaction with gold might have resulted in the fungal cell lysis, and consequent release of intracellular content and DNA damage (**Figure 9D**).^42^

Chi-TY-AuNP’s was found to promote ROS generation in biofilms of both the *Candida* spp. (**Figure 8**). In our study, we hypothesized that various intracellular events mediated the increased levels of ROS production in the Chi-TY-AuNP’s treated *Candida* biofilms. Positively charged Chi-TY-AuNP’s could be easily attracted and adsorbed by the negatively charged cell membrane through electrostatic interaction resulting in pore formation. This leads to the increased permeation and internalization of the Chi-TY-AuNP’s within the mitochondria, followed by the mitochondrial dysfunction and activation of molecular signaling pathways responsible for ROS generation and cell apoptosis (**Figure 9C**).^42^ The findings of Chi-TY-AuNP’s mediated ROS generation in biofilms of both the *Candida* spp. was also evident by the presence of high fluorescence intensity in microscopic fluorescence images (**Figure 8**).

It is known that ergosterol provides stability to the fungal cell wall and maintains its integrity. In addition to intracellular stress, the potential of Chi-TY-AuNP’s in disrupting cell wall integrity of *Candida* spp. was also analyzed in terms of ergosterol content. Furthermore, previous studies reported the synergistic effect of TY and azoles against *Candida* spp. And showed possible interaction of TY with ergosterol.^7^ Thus, the present investigation was extended to estimate ergosterol content in both the *Candida* spp. in response to Chi-TY-AuNP’s. In *C. glabrata* cells, the ergosterol content was significantly decreased in response to Chi-TY-AuNP’s while it remained unchanged in *C. albicans* (**Figure 7C**). This variation in the ergosterol content between *Candida* spp. in response to Chi-TY-AuNP’s might be due to differences in their phylogenetic origin, ploidy, morphology, cell membrane composition and mitochondrial functions.^43^

Although *C. albicans* remains the predominant cause of *Candida*-related infections, over the last decade prevalence of *C. glabrata* mediated infections has considerably increased, resulting in high mortality rate.^3, 44^ Previous studies showed azole like function of exogenously administered tyrosol resulting in inhibition of *Candida* spp. but lacks mechanistic elucidation of its action.^7, 29^ Therefore, it is imperative to address the void in treatment strategies for non-*C. albicans* spp., and asserted to explore the effective therapeutic regimen for *C. glabrata* related infections. Since, in our study Chi-TY-AuNP’s have significantly affected the majority of biochemical parameters (ROS generation, cell surface hydrophobicity, ECM composition and membrane ergosterol content) in *C. glabrata* related to biofilm inhibition for this reason; transcriptional expression analysis of selected genes was studied in *C. glabrata* only. All selected genes of ergosterol biosynthesis, efflux, sterol transporter, and glucan biogenesis were downregulated in response to TY and Chi-TY-AuNP’s except *ERG11* and *CDR1* were upregulated (**Table 1**). The data suggested a direct effect of Chi-TY-AuNP’s on ergosterol, glucan synthesis and efflux pumps of *C. glabrata* and indicated its targeted attack on *C. glabrata* cell membrane, resulting in increased susceptibility against external stress and weak cellular defense leading to biofilm inhibition and eradication (**Figure 9C**).

## CONCLUSIONS

In the present investigation, Chi-TY-AuNP’s were synthesized and characterized via various biophysical techniques. Synthesized Chi-TY-AuNP’s showed fungicidal and biofilm eradication potential against both the *Candida* spp. Further, biochemical studies revealed the interference of Chi-TY-AuNP’s with generated ROS, ECM components and ergosterol content in biofilms of both the *Candida* spp. Transcriptional analysis of selected genes of *C. glabrata* manifested downregulation of genes involved in the maintenance of cell wall biosynthesis. Finally, a limitation of this study is that we performed the transcriptional analysis of *C. glabrata* only and further research is warranted to elucidate the therapeutic and mechanistic effect of Chi-TY-AuNP’s against different biofilm-forming pathogens. Our findings confirm the effectiveness of an alternative therapeutic system that may control *Candida*-associated infections. From the above study we may conclude that Chi-TY-AuNP’s may act effectively in biofilm eradication when applied/ coated on clinically relevant biomaterials and eventually help in will help in preventing implant-associated fungal infections.

Future research needs to be comprehensively focused on the critical evaluation of cytotoxicity, biodistribution, pharmacodynamics and pharmacokinetics of such nanocarrier systems while investigating their efficacy against biofilm-associated infections.

## ABBREVIATIONS

Chi-TY-AuNP’s: Tyrosol functionalized chitosan gold nanoparticles
eDNA: Extracellular DNA
ECM: extracellular matrix
QS: quorum sensing
QSM: quorum sensing molecule
TY: tyrosol
MDR: multidrug resistance
ROS: reactive oxygen species
EPR: enhanced permeability and retention
SPR: surface plasmon resonance
PDI: polydispersity index
DLS: dynamic light scattering
AFM: atomic force microscopy
FTIR: fourier-transform infrared
HRTEM: high-resolution transmission electron microscopy
IR: infra-red
DLE: drug loading efficiency
YPD: yeast extract peptone dextrose agar
RPMI: rosewell park memorial institute
MTP: 96-well multitier plate
MIC: minimum inhibitory concentration
MFC: minimum fungicidal concentration
BIC_80_: biofilm inhibitory concentrations
BEC_80_: biofilm eradication concentration
FESEM: field emission scanning electron microscopy
FDA: fluorescein diacetate
PI: propidium iodide
DCFDA: 2,7-dichlorodihydrofluoroscein diacetate
PCI: isoamyl alcohol
HI: hydrophobicity index
CT: cycle threshold
SD: standard deviation
SAED: selected area electron diffraction

## ACKNOWLEDGMENTS

TCY acknowledges Ministry of Human Resource Development (MHRD), Government of India, for financial support as a Senior Research Fellowship. PG acknowledges the DBT-RAship program, Department of Biotechnology, Government of India. The authors are grateful to Institute Instrumentation Centre, IIT Roorkee, Uttarakhand, India, for providing the instrumentation facilities.

## AUTHOR CONTRIBUTIONS

TCY and PG equally contributed to this work. All authors contributed to data analysis and manuscript writing. All authors have approved the final version of the manuscript.

---

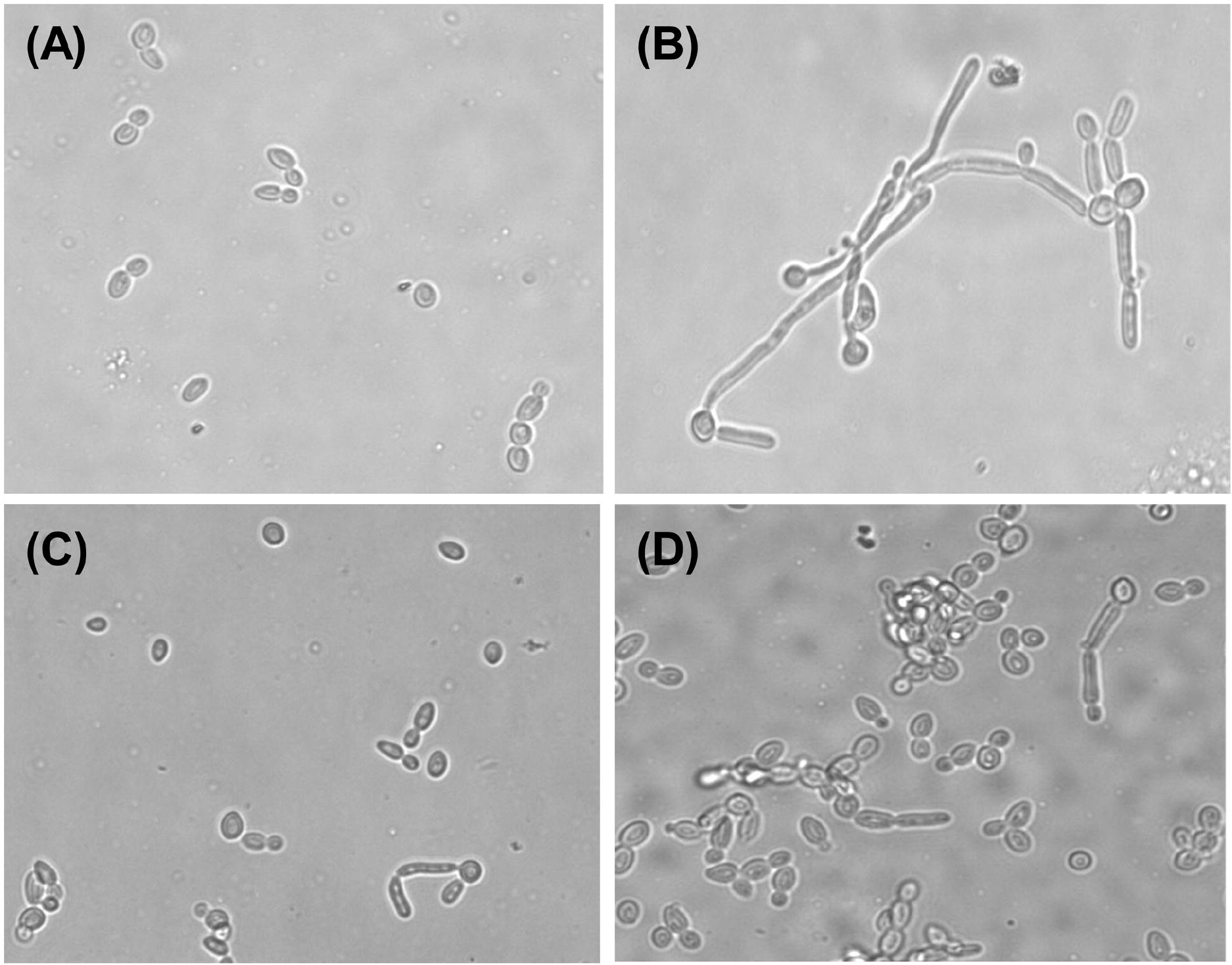

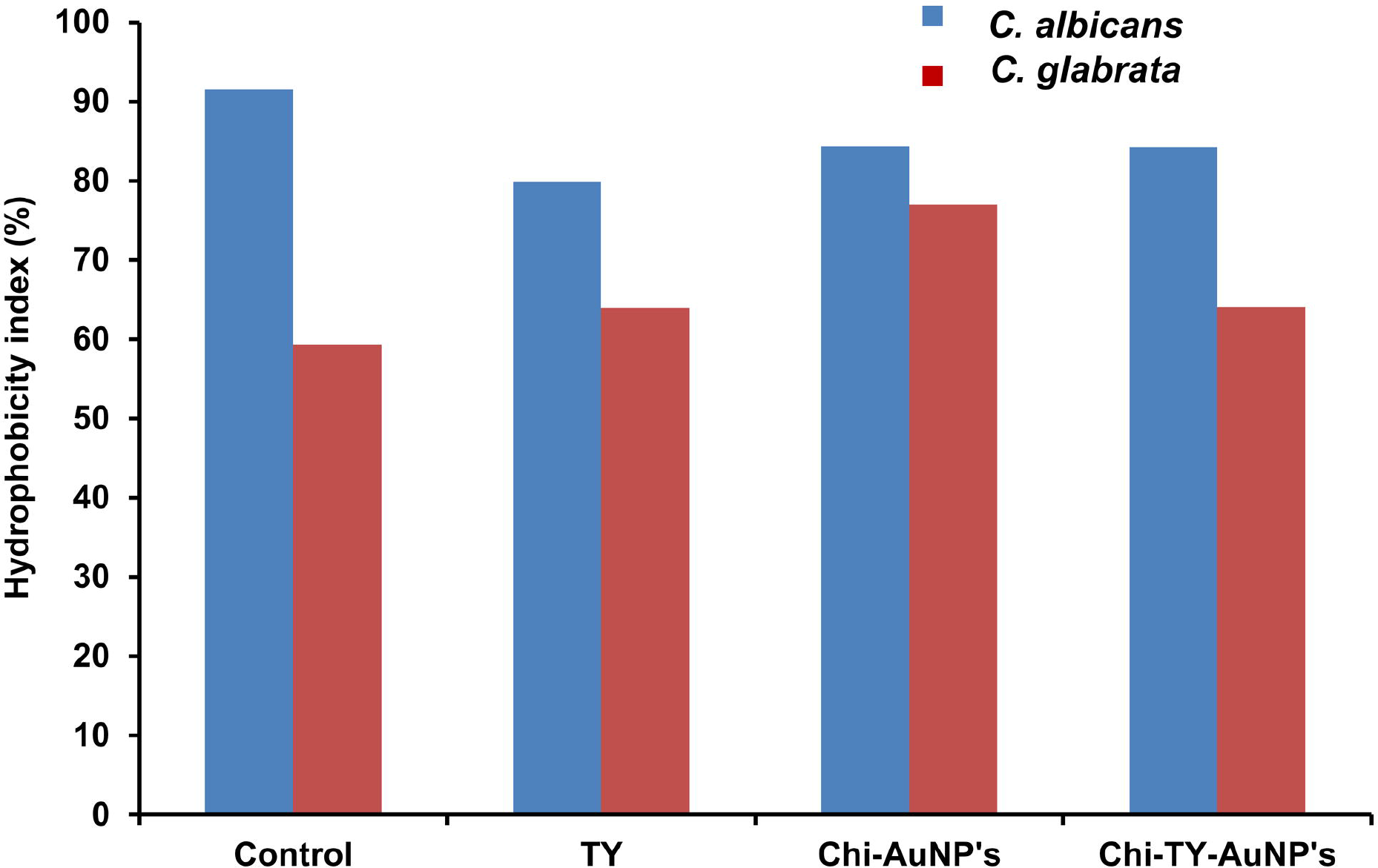

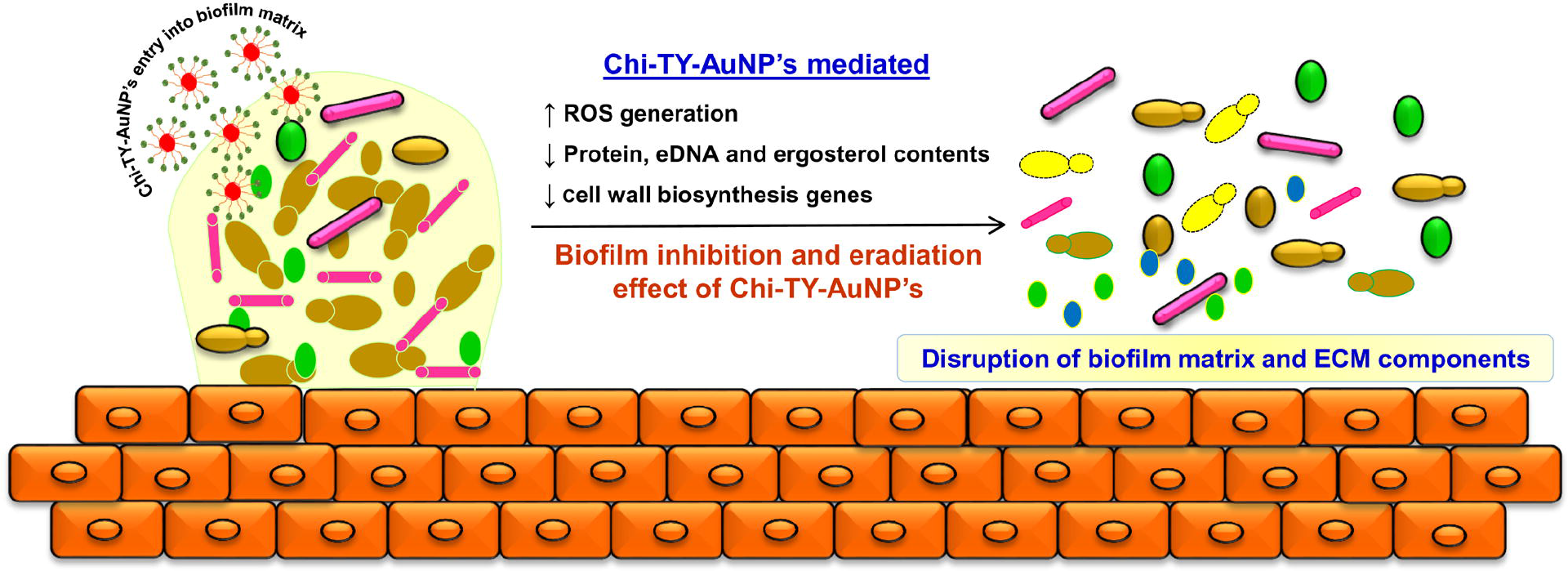

