## Supplemental Table for "Plausible Mechanistic Insights in Biofilm Eradication Potential of against *Candida* spp. using In Situ Synthesized Tyrosol Functionalized Chitosan Gold Nanoparticles as a versatile Antifouling Coating on Implant Surfaces" (from another version of this preprint: version 1, 2021-10-01)

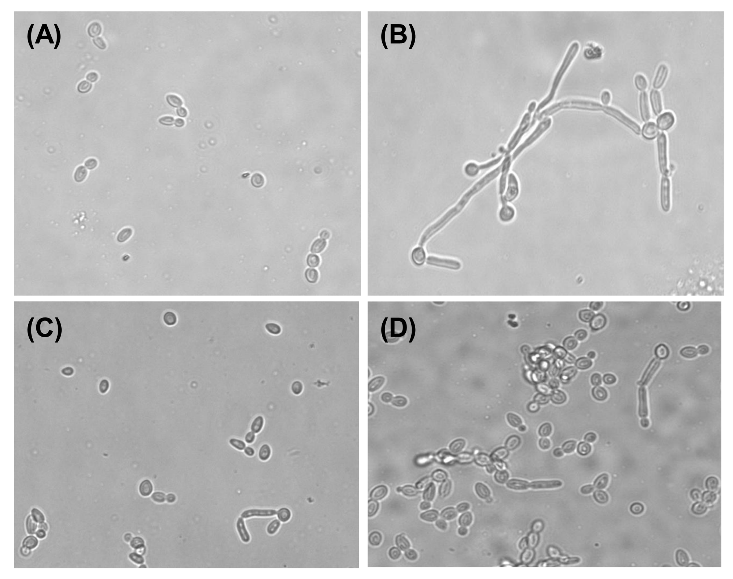

**Figure S1.** Inhibition of germ tube formation in *C. albicans* (A) untreated or negative control cells (B)10% FBS treated cells or positive control cells (C) Chi-TY-AuNP's treated cells (D) Chi-AuNP's treated cells.

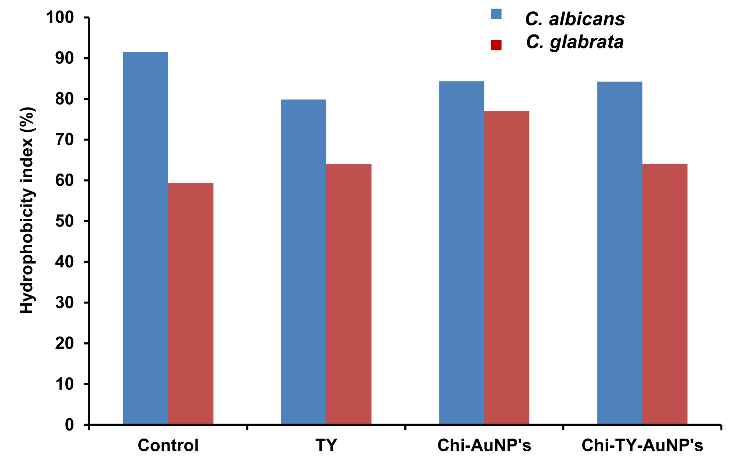

**Figure S2.** Quantification of hydrophobicity index (HI) in *C. albicans* and *C. glabrata* cells treated with subinhibitory concentrations of TY, Chi-AuNP's and Chi-TY-AuNP's.
